# Early sex-specific organ transcriptional divergence without physiological differences in a murine model of fecal-induced peritonitis

**DOI:** 10.64898/2026.02.18.706550

**Authors:** Alexandra Troitskaya, Aminmohamed Manji, Onon Batnyam, Eric K. Patterson, Forough Jahandideh, Dhruva J. Dwivedi, Manoj M. Lalu, Asher A. Mendelson, Braedon McDonald, Stephane L. Bourque, Kimberly F. Macala, Alison E. Fox-Robichaud, Patricia C. Liaw, Gediminas Cepinskas, Ruud A. W. Veldhuizen, Sean E. Gill, the National Preclinical Sepsis Platform, The Canadian Critical Care Translational Biology Group, and Sepsis Canada

## Abstract

Sepsis is defined as a dysregulated response to infection, leading to life-threatening organ dysfunction that particularly affects parenchymal organs. Clinical studies remain inconclusive regarding the impact of biological sex on sepsis, and preclinical studies are predominantly performed in male animals. We examined early (8 h) septic responses in male and female mice using a fecal-induced peritonitis (FIP) model. Blood biochemical parameters, body temperature, and murine sepsis scores provided evidence of a septic response in animals randomized to FIP compared to controls, but showed no physiological differences between male and female mice. Transcriptomic analysis of the liver, kidney, and lung showed consistent inflammatory activation in response to sepsis as compared to controls. Notably, in the kidney and lung, female mice exhibited stronger immune activation and a heightened inflammatory response compared to males. Thus, biological sex differences in the septic response can be detected in early acute sepsis without apparent physiological differences.

## Introduction

Sepsis, which is defined as a life-threatening organ dysfunction caused by a dysregulated host response to infection, is a global health concern with 48.9 million cases and 11 million deaths annually [1–5]. The pathophysiology of sepsis includes dysfunction of multiple organs with significant involvement of the lungs, liver, and kidneys [6–8]. Further, sepsis occurs in both male and female patients with various reports suggesting differences in incidence, ICU admissions, and mortality [9–12]. To date, there is no effective pharmacological therapy for sepsis, likely due to the large variety of mechanisms driving sepsis pathophysiology in multiple organs, in both male and female patients [13–15]. Research taking a wide-ranging approach to investigate pathophysiological pathways is required to better understand the development of sepsis.

One approach to analyze the broad molecular patterns associated with sepsis is via one or more of the existing “*omics*” tools, such as metabolomics, proteomics, and transcriptomics. This approach has been employed with clinical samples to stratify sepsis patients into molecular endotypes linked to immune response profiles and mortality risk, thereby opening the possibility for precision medicine approaches [16–18]. In particular, transcriptomic profiling via RNA sequencing (RNAseq) has been suggested as a promising tool for early patient stratification, as it captures dynamic molecular signatures prior to the full onset of organ dysfunction [15]. Despite this progress, these clinical studies are limited in the type of samples that can be analyzed, and for a detailed understanding of organ responses, preclinical animal studies are required.

Preclinical animal studies on sepsis vary in both sepsis induction and species, and have been predominantly performed in male animals [19,20]. Although these approaches have been instrumental in advancing our understanding of disease pathophysiology, different models often provide different results [21,22]. To improve translatability of sepsis studies, the National Preclinical Sepsis Platform (NPSP), which is associated with Sepsis Canada, developed a set of rigorous and transparent experimental mouse protocols [23]. This methodology includes a reproducible septic stimulus via fecal-induced peritonitis (FIP), a scoring system to assess physiological sign of sepsis, utilization of both male and female mice, and a biobanking protocol for multiple tissues [24,25].

The goal of this exploratory study was to perform transcriptomic analyses of multiple organs in mixed biological sex population of mice using the NPSP-standardized mouse sepsis protocol. Additionally, we explored biological sex differences in these responses in both the kidney and lung.

## Materials & Methods

### Animals

This blinded study used 10–12-week-old C57BL/6 male (24.4 ± 1.2 g) and female (20.3 ± 1.4 g) mice purchased from Charles River Laboratories. Mice were randomized into four groups: male control (n = 3), male septic (n = 4), female control (n = 3), and female septic (n = 5). Mice were evaluated based on total body weight, body temperature and the modified murine sepsis score (MSS) criteria, which included five parameters: appearance, posture, activity, response to stimulus, and respiration quality. Each parameter was scored from 0 (normal) to 3 (severe). Assessments were performed by qualified personnel (2-3 people) who were blinded to experimental group, and average scores were recorded at 0, 4, and 8 h post-injection to monitor the animals’ response to sepsis and to ensure humane endpoints were met. Criteria for early euthanasia included any of the following: loss of body temperature or body weight (≥ 20% baseline), an MSS score of 3 in any of the categories, or a total MSS score ≥ 9.

At 0 h, mice were assessed, anesthetized with isoflurane, and given an intraperitoneal injection of 0.75 mg/g body weight rat fecal slurry suspended in 5% dextrose/10% glycerol (septic) or 5% dextrose/10% glycerol alone (control). At 4 h, mice were again assessed and then given a subcutaneous injection of 0.1 mg/mL Buprenorphine in Ringers Lactate at 0.05 mg/kg body weight. At 8 h, mice were once again assessed and following anesthesia with isoflurane, peritoneal lavage fluid (PLF) was collected via phosphate-buffered saline (PBS) lavage. Mice were then exsanguinated by carotid cutdown, and arterial blood was collected for analysis using the epoc® Blood Analysis System. PLF samples were plated on blood agar and incubated for 24 h to assess bacterial growth. Liver, lung, and kidney tissues were harvested, snap-frozen with liquid nitrogen, and stored at −80°C. The animal procedures followed the NPSP harmonized SOPs (SOP-NPSP-MTD-01-04); see NPSP-01 protocol for full methodological details [24]. All animal procedures were conducted in accordance with the Canadian Council on Animal Care guidelines and approved by the Western University Animal Care Committee (AUP #2022-023).

### Illumina NovaSeq Next Generation Sequencing

Liver, kidney, and lung tissue RNA samples were extracted and sequenced at Génome Québec (Genome Canada, Montréal, Québec) using the Illumina NovaSeq PE100 (Illumina Inc., San Diego, CA). Total RNA was extracted using the RNeasy Plus Universal Kit from QIAGEN (cat. 73404) with a homogenization step using QIAGEN TissueLyser II with 5 mm stainless steel beads (cat. 69989). The extraction was performed according to the manufacturer’s instructions. Total RNA was quantified, and its integrity was assessed with LabChip GXII (PerkinElmer) instrument. The minimum cutoff for RNA quality was an RNA integrity number ≥ 3.9; however, the average RNA integrity number of the samples used was 7.0. Libraries were generated from 100 ng of total RNA. mRNA enrichment was performed using the NEBNext Poly(A) Magnetic Isolation Module (New England Labs). cDNA synthesis was achieved with NEBNext UltraExpress RNA Library Prep Kit (New England BioLabs), as per the manufacturer’s recommendations.

The library was sequenced on the Illumina NovaSeq 6000 (200 cycles) as paired end reads (2 × 100 bp), with approximately 3200M reads per run (∼200M reads per sample). A phiX library was used as a control and mixed with libraries at the 1% level. Program BCL Convert 4.2.4 was then used to demultiplex samples and generate fastq reads. Fastq data files were analyzed using Partek Flow (version M-GL-03002 v1.0; St. Louis, MO; https://www.illumina.com/products/by-type/informatics-products/partek-flow.html). After importation, the data were aligned to the *Mus musculus* genome mm39 using STAR 2.7.8a and annotated using Ensembl Transcripts 112. Gene features with more than three mapped reads were normalized using the DESeq2 median ratio method with a noise reduction filter. Principal component analysis (PCA) was generated using variance-stabilized counts to visualize overall sample clustering and assess sources of variation across experimental conditions. Volcano plots were generated based on DESeq2 (FC ≥ 1.5 FDR ≤ 0.05) and then plotted with RStudio using the packages *ggplot2* and *dplyr*. For male vs. female comparisons, the DESeq2 lists were entered into Venny 2.1 (https://bioinfogp.cnb.csic.es/tools/venny/) to find the number of shared and exclusive differentially expressed genes (DEGs) in male vs. female. The annotated volcano plots with top 30 inflammatory markers were created in RStudio using additional packages *biomaRt* and *ggrepel*. The selection of markers was based on the list of 120 conventional inflammatory markers in mice curated using ChatGPT based on the most expressed genes in mouse sepsis studies [26–30] and pathways from the following databases: MSigDB [31], InnateDB [32], and KEGG [33]. The top expressed inflammatory markers in each sex were visualized in RStudio as a heatmap using packages *dplyr*, *tibble*, *biomaRt*, *ComplexHeatmap*, *circlize*, and *grid*.

### Gene Ontology (GO) Analysis

Separate lists of upregulated and downregulated genes were generated for each tissue for combined population organ response comparisons. For each comparison, gene lists were filtered using DESeq2 (FC ≥ 1.5; FDR ≤ 0.05) and submitted to Metascape (https://metascape.org/gp/index.html) using express analysis of *Mus musculus* gene IDs [34]. For combined population response in liver, kidney, and lung, the top 20 mapped GO pathways were put through Venny 2.1 to find pathways that were present across all three tissues. The pathways were then plotted with RStudio using packages *dplyr*, *tidyr*, and *ggplot2*.

### Gene Set Enrichment Analysis (GSEA)

All genes that passed quality control in all the assigned comparisons (combined population and separate biological sexes in different organs) were subjected to the GSEA software (version 4.4.0; Broad Institute; http://www.gsea-msigdb.org/gsea/index.jsp) for pathway enrichment analysis [35,36]. GSEA was performed with the default settings. Briefly, the threshold per pathway ranged from a minimum of 15 to a maximum of 500 genes. Permutation was conducted 1,000 times following the default weighted enrichment statistics and signal-to-noise metric for ranking genes according to differential expression levels across two compared phenotypes. The hallmark gene set collection (v2025.1.Mm) of mouse-ortholog pathways from Molecular Signatures Database was used as an enrichment database (MSigDB orthology-mapped 50 gene sets; mouse Ensembl Gene ID platform; https://www.gsea-msigdb.org/gsea/msigdb/index.jsp). Visualization was performed using RStudio using the packages *ggplot2*, *dplyr*, *scales*, and *viridis*. The significant gene sets were defined as those with an FDR ≤ 0.1. The 50 hallmark pathways were split into 6 categories: Immune & Inflammation, Stress & Damage Response, Proliferation & Cell Cycle, Metabolism, Canonical Signalling, and Development & Differentiation.

### Statistical Analysis

Data analysis and visualization were performed using GraphPad Prism (version 9.4.1; GraphPad Software Inc., San Diego, CA; https://www.graphpad.com/features). Results are presented as means ± standard error of the mean (SEM). For body temperature and MSS total score, a three-way ANOVA (α = 0.05) was used to assess the effects of time, biological sex, and sepsis on the measured outcomes. Tukey’s multiple comparisons test (p ≤ 0.05) was applied for post hoc pairwise comparisons. For PLF and blood measurements, a two-way ANOVA was used with sepsis status, biological sex, and their interactions. Tukey’s multiple comparisons test (p ≤ 0.05) was applied for pairwise comparisons.

## Results

All animals in our study survived to the 8 h post-injection period and with similar physiological responses observed due to slurry (septic) compared to dextrose (control) in male and female animals (Fig 1). Both male and female animals exposed to slurry exhibited a significant reduction in core body temperature (Fig 1A) and a significant increase in MSS over time compared to animals receiving dextrose (Fig 1B). Additionally, PLF cultures exhibited significantly higher bacterial counts in slurry animals relative to dextrose controls (Fig 1C) in both sexes. Blood analysis showed significant metabolic disturbances in the slurry group, with glucose, sodium, and pH levels significantly reduced, and blood urea nitrogen (BUN) levels significantly elevated compared with the dextrose group (Fig 1D) with no statistically significant differences observed between males and females.

**Fig 1.**
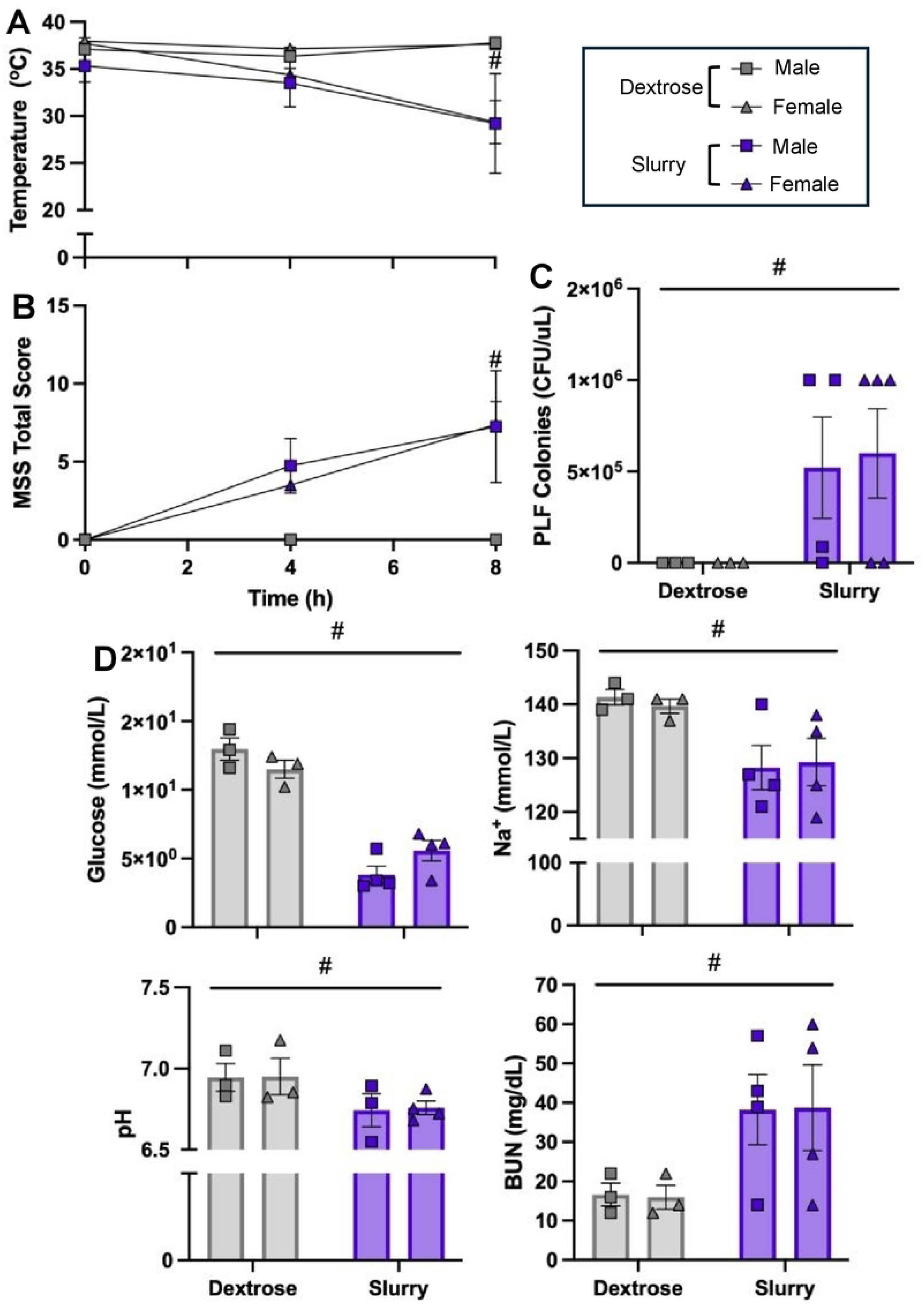
Physiological Response to Acute Sepsis in Male and Female Mice. **(A)** Body temperature at 0, 4, and 8 h, showed significant declines in mice that received slurry (septic) at 4 and 8 h. **(B)** MSS scores at 0, 4, and 8 h, revealed significant increases in mice that received slury (septic) at 4 and 8 h. **(C)** Bacterial counts in PLF samples were significantly higher in slurry vs. dextrose mice. **(D)** Blood measurements of glucose, sodium, pH, and BUN: glucose, sodium, and pH decreased, while BUN increased in slurry (septic) mice. Data are presented as mean ± SEM. Statistical analyses were performed by three-way ANOVA (A–B) or two-way ANOVA (C–D) with Tukey’s post hoc test (# p ≤ 0.05, * p ≤ 0.05). Sample sizes: dextrose (control: male n = 3; female n = 3) and slurry (septic: male n = 4; female n = 5).

To evaluate the general organ transcriptomic responses to slurry, combined male/female cohorts of liver, kidney, and lung samples were analyzed. As shown in Fig 2, slurry exposure elicited extensive transcriptional changes in all three organs compared to dextrose exposure. PCA demonstrated a clear separation between slurry and dextrose samples, and this separation appeared to be most notable in the liver (Fig 2A), compared to transcriptional responses in the kidney and lung (Fig 2B-C). Analysis of the DEGs in each tissue via volcano plots revealed a number of up- and downregulated genes in each tissue due to slurry exposure (Fig 2D-F). The top 20 most significant GO pathways associated with the combined up- and downregulated DEGs in each organ revealed remarkable conservation between organs with 13 pathways shared among the three different organs (Fig 2G-I). Many of these pathways were associated with inflammation and dominated by upregulated DEGs. Five pathways, all present within the lung, were shared with either kidney and liver and were generally also associated with immune function and inflammation. Further, 11 pathways were not shared among the top 20 pathways of each organ.

**Fig 2.**
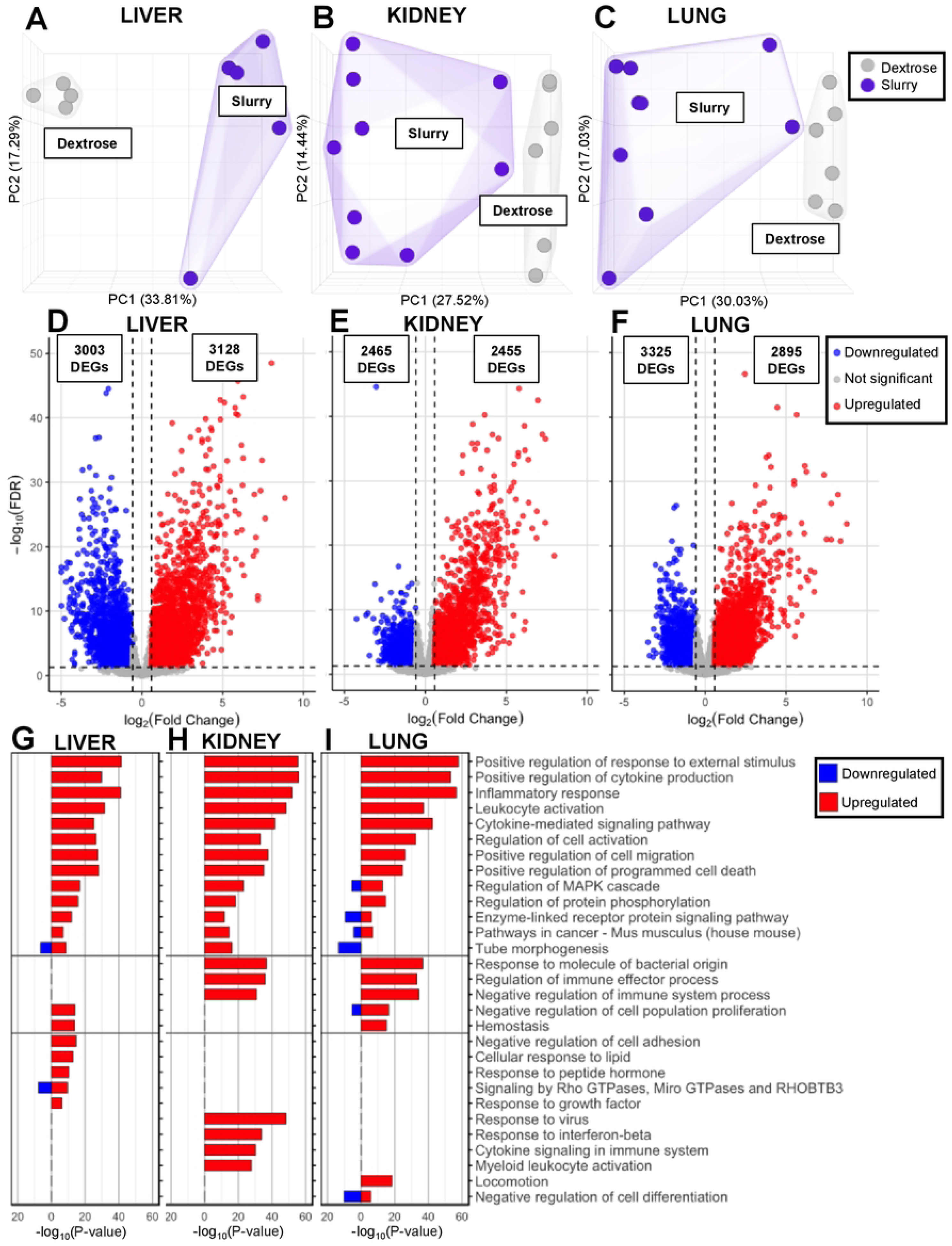
Liver, Kidney, and Lung Transcriptional Septic Responses in the Combined Population. **(A-C)** Liver, kidney, and lung PCA each had two distinct clusters based on the dextrose and slurry groups across all tissues. The liver tissue exhibited the clearest separation amongst the two groups. **(D)** Liver DEGs: downregulated = 3,003 DEGs; upregulated = 3,128 DEGs. **(E)** Kidney DEGs: downregulated = 2,465 DEGs; upregulated = 2,455 DEGs. **(F)** Lung DEGs: downregulated = 3,325 DEGs; upregulated = 2,895 DEGs. **(G-I)** The GO in all three tissues was significantly upregulated in immune response and inflammatory activation. The significance is indicated as -log_10_(P) by bar length. Sample sizes: liver dextrose (control, n = 4), slurry (septic, n = 5); kidney and lung dextrose (control, n = 6), slurry (septic, n = 9).

To complement the DEGs and GO analysis, we performed gene set enrichment analysis (GSEA), which does not rely solely on DEGs. GSEA analysis was performed with 50 available hallmark pathways reflecting well-defined biological processes that, for ease of interpretation, were subdivided into 6 categories (Fig 3). GSEA demonstrated statistically significant enrichment of the Immune & Inflammation and Stress & Damage Response pathways in the slurry (septic) vs. dextrose (control) group across all three tissues (Fig 3A-B). Proliferation & Cell Cycle pathways were also mostly enriched in mice from the slurry group in all three tissues (Fig 3C). The Metabolism and Canonical Signalling pathways had a very similar enrichment pattern in the kidney and lung, with most pathways enriched in response to slurry; however, within the liver, only a small portion of pathways became enriched with slurry, while others were enriched in the dextrose group (Fig 3D-E). Lastly, the pathways associated with Development & Differentiation had the most variability of enrichment across all three tissues, with the majority of pathways not reaching significant enrichment in either experimental group (Fig 3F).

**Fig 3.**
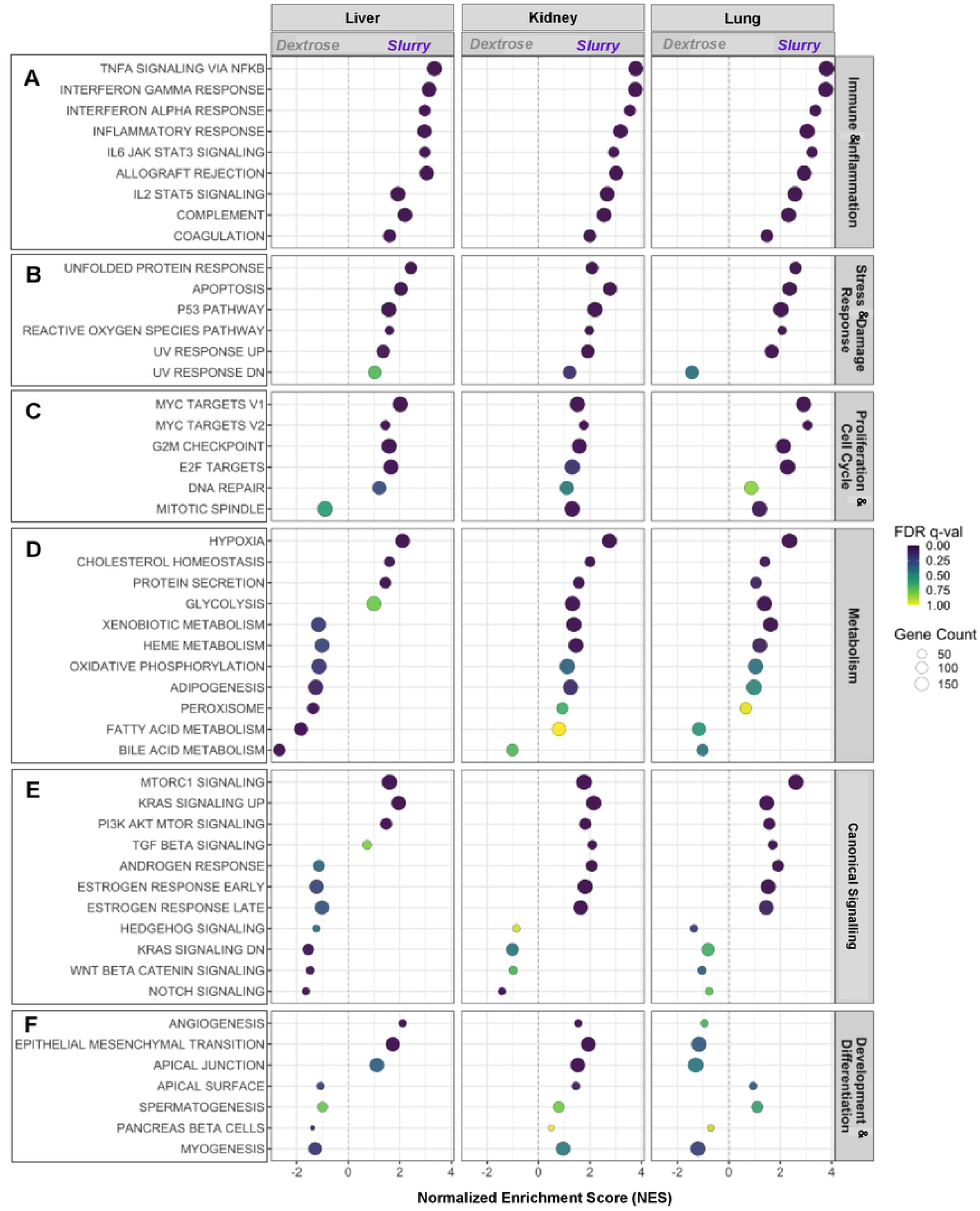
Liver, Kidney, and Lung Transcriptional Septic Responses in the Combined Population: GSEA using the 50 Hallmark Pathways. The 50 hallmark pathways were split into 6 categories: **(A)** Immune & Inflammation, **(B)** Stress & Damage Response, **(C)** Proliferation & Cell Cycle, **(D)** Metabolism, **(E)** Canonical Signalling, and **(F)** Development & Differentiation. **(A)** All Immune & Inflammation, **(B)** Stress & Damage Response, and **(C)** Proliferation & Cell Cycle pathways were collectively enriched in the slurry (septic) group for all three tissues with minor variations. **(D)** The pattern of enrichment for Metabolism pathways was similar across kidney and lung, with enrichment in the slurry (septic) group; in liver, more pathways were enriched in the dextrose (control) group. **(E)** Canonical Signalling was primarily enriched in kidney and lung in the slurry (septic) group; in liver, the enrichment appeared to be evenly distributed between the two groups. **(F)** Development & Differentiation enrichment appeared to be varied in each tissue, with liver and kidney having enrichment in slurry (septic) group. No Development & Differentiation pathways were significantly enriched within the lung tissue. The cutoff for significance is FDR ≤ 0.1. The normalized enrichment score (NES) reflects the direction and magnitude of pathway enrichment in females relative to males. The gene count per pathway is identified by the size of the circle.

A similar strategy of exploring DEGs and GSEA analysis was performed to dissect potential sex-based differences in responses, focusing on the inflammation and injury-associated pathways, as these represented the most robust signals at the whole organ level. This analysis of sex-stratified transcriptional responses, however, was only performed in the kidney and lung due to reduced n-values and subsequently, a lack of sufficient power within the liver samples after quality control assessment of the transcriptomic analysis.

Within the kidneys, a total of 1,892 DEGs were shared between males and females in response to slurry, while 516 DEGs and 2,310 DEGs were observed exclusively in males and exclusively in females, respectively (Fig 4A). The majority of DEGs appeared to be upregulated in response sepsis in both males and females (Fig 4B-C). Interestingly, while comparison of inflammatory genes by heatmap showed a general upregulation of inflammatory genes in the slurry vs. dextrose groups, there appeared to be a more pronounced upregulation in the female animals (Fig 4D). GSEA also revealed sex differences in pathway enrichment patterns in both dextrose and slurry conditions (Fig 5). Specifically, in both the dextrose and slurry groups, most of the Immune & Inflammation pathways as well as the Stress & Damage Response pathways were upregulated in female compared to male mice (Fig 5A-B). Differences within the kidney between males and females in other hallmark pathways revealed limited sex-based differences in the dextrose groups that were generally maintained in the slurry groups (Sup Fig S1).

**Fig 4.**
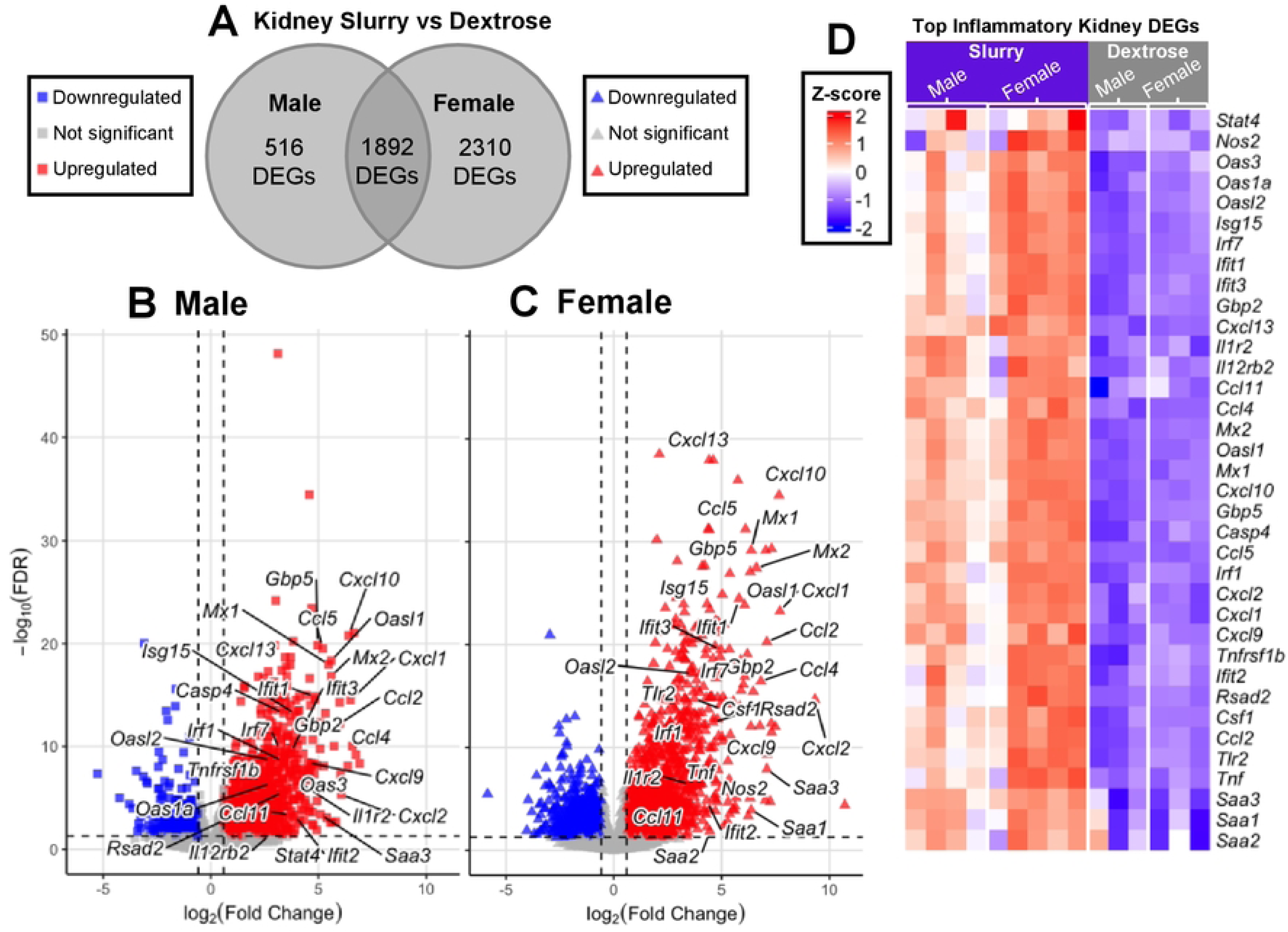
The Effect of Biological Sex on the Transcriptional Response of the Kidney During Sepsis. **(A)** In mice receiving slurry (septic), there were gene transcripts that were shared by both sexes as well as many that were exclusive to one sex or the other in the kidney: 1,892 DEGs shared, 516 DEGs exclusively in males, and 2,310 DEGs exclusively in females. **(B-C)** Volcano plots illustrate the variability in fold change and significance in DEGs in the kidney in male vs. female mice in reponse to slurry (septic). **(D)** Heatmap represents relative gene expression of hightly upregulated inflammatory genes in male and female mice under dextrose (control) and slurry (septic) conditions (blue – downregulated; red – upregulated). Male and female dextrose groups had consistent downregulation of inflammatory genes. Animals that received slurry (septic) exhibited an overall upregulation of the inflammatory genes. This increase appeared to be augmented in females compared to males.

**Fig 5.**
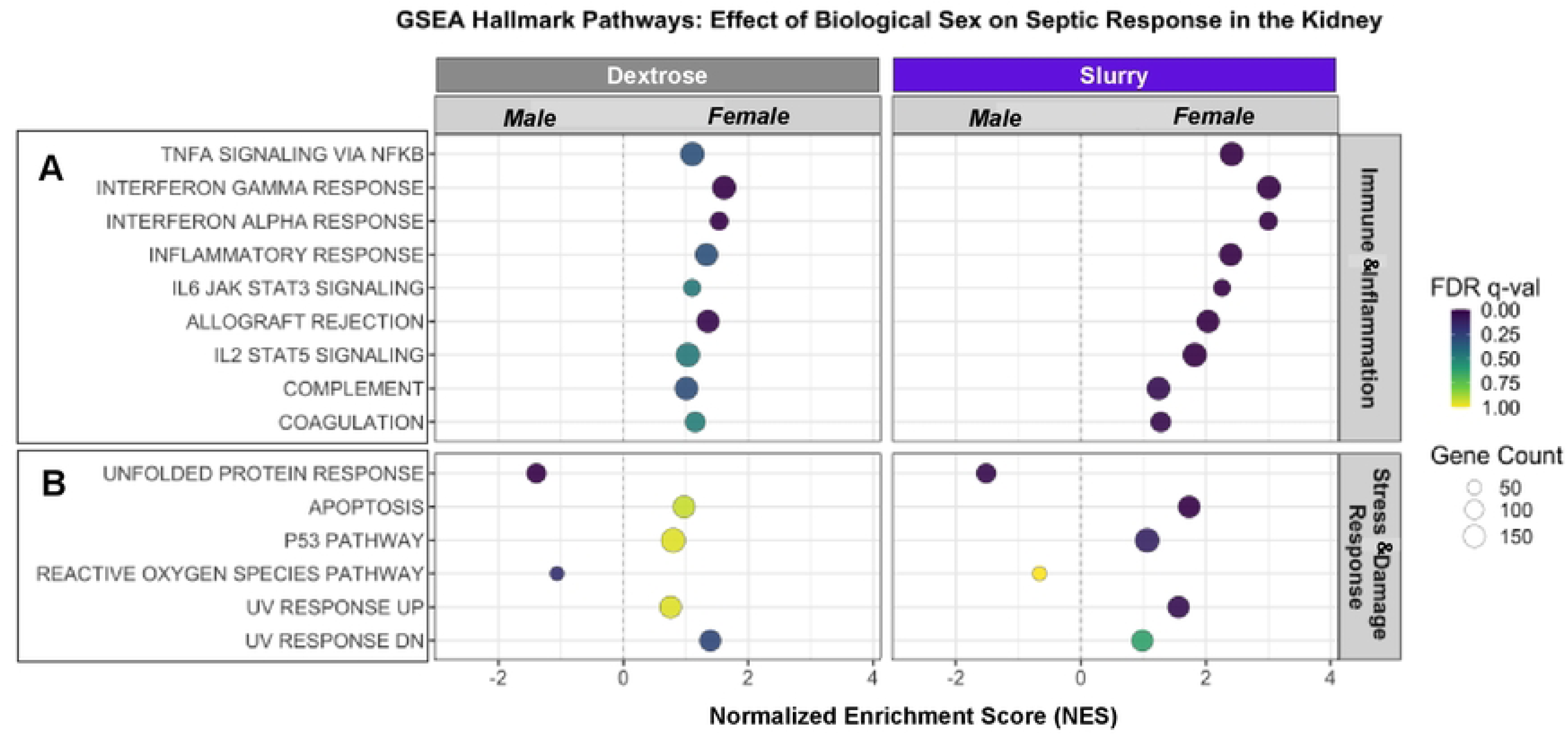
GSEA in the Kidney during Sepsis in Male vs. Female Mice. Females that received slurry (septic) had a significant enrichment in all **(A)** Immune & Inflammation pathways as well as some **(B)** Stress & Damage Response pathways. The cutoff for significance is FDR ≤ 0.1. The normalized enrichment score (NES) reflects the direction and magnitude of pathway enrichment in females relative to males. The gene count per pathway is identified by the size of the circle.

Within the lung, sex-based analyses of the transcriptional responses revealed similar findings as those observed within the kidney (Fig 6-7). Among the genes expressed during sepsis, 2,447 DEGs were shared between sexes, while 682 DEGs and 2,407 DEGs were observed only in males and only in females, respectively (Fig 6A). As observed in the kidneys, there was a consistent upregulation of inflammatory genes in animals in the slurry vs. dextrose groups with an augmented upregulation observed in the female animals (Fig 6B-D). GSEA analysis revealed that in the dextrose animals, males exhibited enrichment across all displayed pathways (Fig 7). However, in the slurry group (sepsis), this pattern shifted towards the female animals being enriched in most Immune & Inflammation and Stress & Damage response pathways (Fig 7A-B). Outside of the inflammatory and tissue damage pathways, the sex-based differences within the lung for other hallmark pathways appeared to have a varied pattern of responses (Sup Fig S2).

**Fig 6.**
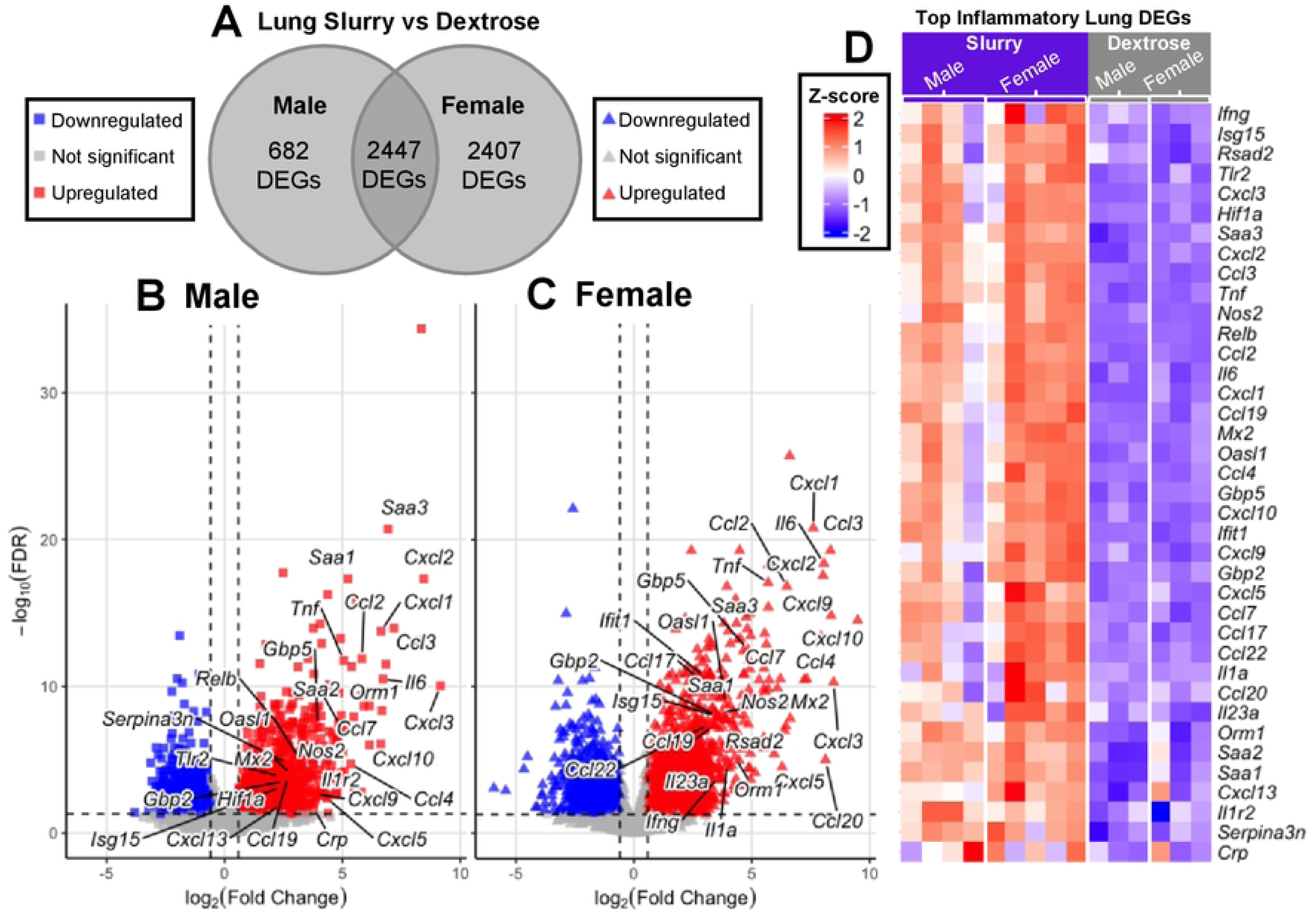
The Effect of Biological Sex on the Transcriptional Response of the Lung During Sepsis. **(A)** In mice receiving slurry (septic), there were gene transcripts within the lung that were shared by both sexes as well as many that were exclusive to either males or females: 2,447 DEGs shared, 682 DEGs exclusively in males, and 2,407 DEGs exclusively in females. **(B-C)** Volcano plots illustrate the variability in fold change and significance in DEGs in the lung in male vs. female mice in reponse to slurry (sepsis). **(D)** Heatmap represents relative gene expression of the most upregulated inflammatory genes in male and female mice under dextrose (control) and slurry (septic) conditions (blue – downregulated; red – upregulated). Male and female dextrose groups had consistent downregulation. Animals that received slurry exhibited an overall upregulation in the inflammatory response, and females appeared to have an overall stronger inflammatory signal, compared to males.

**Fig 7.**
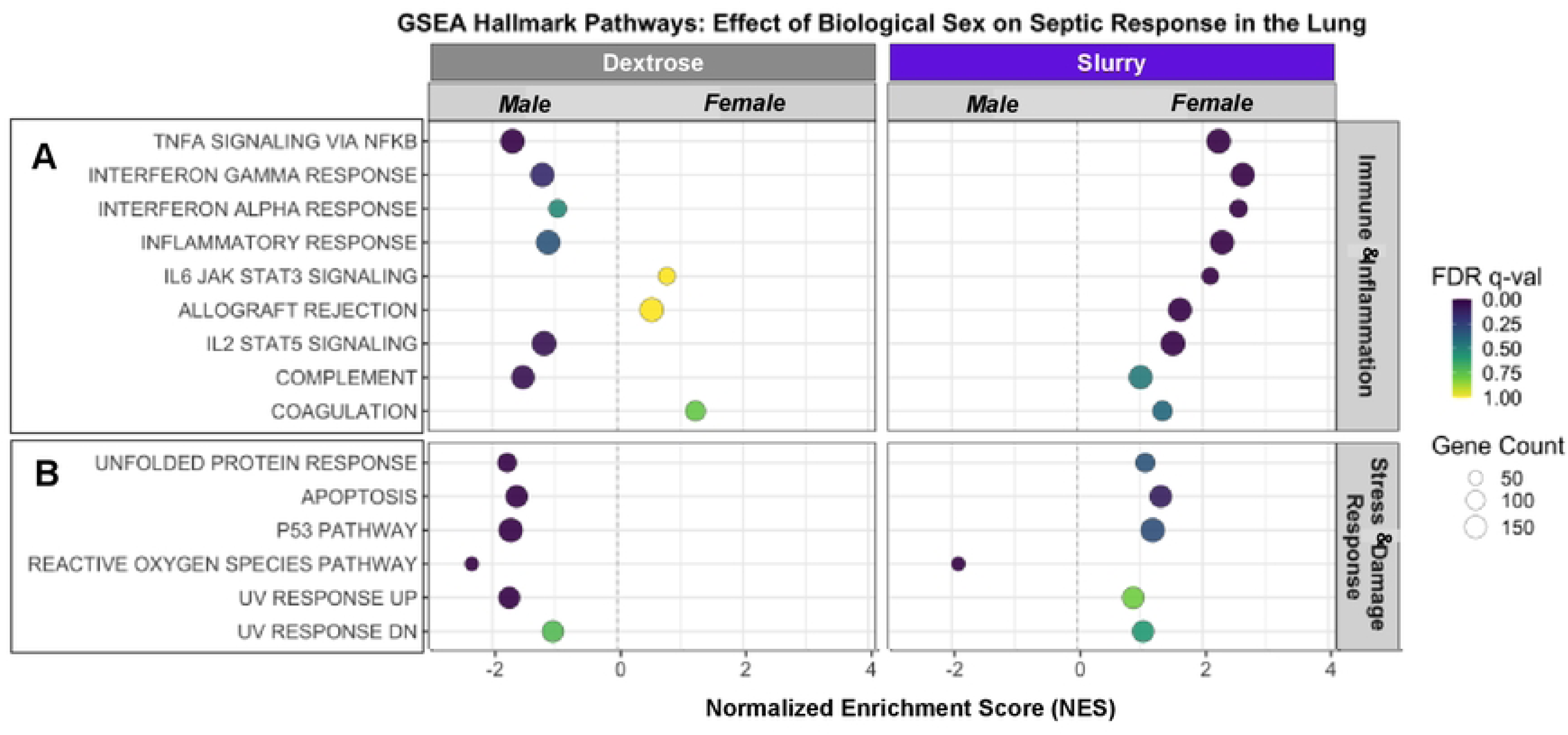
GSEA in the Lung during Sepsis in Male vs. Female Mice. Males exhibited enrichment of most **(A)** Immune & Inflammation and **(B)** Stress & Damage Response pathways under dextrose (control) conditions, whereas females displayed enrichment of these pathways under slurry (septic) conditions. The cutoff for significance is FDR ≤ 0.1. The normalized enrichment score (NES) reflects the direction and magnitude of pathway enrichment in females relative to males. The gene count per pathway is identified by the size of the circle.

## Discussion

Based on the clinical impact of sepsis and the limitations in animal models, such as various experimental models, lack of sex-stratified research, and limited broad organ assessments, this exploratory study examined the transcriptional responses within the liver, kidney, and lung in male and female mice in an established preclinical model of sepsis. Our findings verified the development of sepsis in animals receiving an injection of fecal slurry, but not those receiving dextrose, via the MSS scores as well as other parameters. The subsequent transcriptomic analysis revealed significant inflammatory pathway activation occurring in all three organs following sepsis, with the majority of the top 20 enriched pathways overlapping among the three organs. Furthermore, in the kidney and lung, activation of this response was more pronounced in female animals compared to male animals. It is concluded that, although similar pathways are activated in different organs, the extent of this response might be significantly affected by biological sex.

As with any preclinical study, it is important to interpret the data in the context of the experimental design. In the current study, sepsis was induced by injection of a slurry of rat-derived fecal material based on the individual animal’s weight. Previous reviews have emphasized the heterogeneity introduced by different animal models, each with its limitations and variations in pathogen load, host immune responses, and timing [20,21,37]. For example, our model contrasts with other models, such as the commonly used injection of lipopolysaccharide, in which responses are very specific to the activation of a limited number of pathways, such as those associated with Toll-like receptor 4 [38]. Our model is also different from the well-established sepsis model of cecal ligation and perforation (CLP). In the CLP model, the septic insult is the animal’s own fecal material and thus differences in the composition of the animal’s fecal microbiome would result in animals receiving different septic insults. This becomes especially important when examining male and female differences, as they are known to have substantially different microbiomes potentially impacting sepsis progression following CLP [39,40]. Another difference between our approach and the CLP model is that dosing is more difficult to control in the CLP model, which requires surgery, perforation of the cecum, and the subsequent extrusion of fecal material from the cecum. The intraperitoneal injection of fecal slurry provides a more consistent dosing. Interestingly, the normalization of the fecal slurry injection by weight in our model led to our slightly lighter female mice receiving a smaller amount of slurry. This lower amount of slurry given to female mice suggests that the enhanced inflammatory signalling observed in the female kidney and lung is a biological response and not due to the dose. Overall, our data demonstrate organ-specific responses and sex-dependent differences after a standardized polymicrobial insult.

A second important feature of our experimental design is that the assessments were performed at 8 h post-injection of either dextrose or slurry, which reflects a very acute time point in the septic response. At this time, both male and female mice receiving slurry exhibited equivalent physiological responses – including hypothermia, elevated MSS, bacterial burden, and blood biomarker changes, yet sex-based differences were only observed in our transcriptomic analysis. The physiologic observations align with those in other murine models of early sepsis that emphasize similarities in vital signs and systemic indicators at 6 – 12h post-septic insult, capturing acute phase responses [19]. Furthermore, the sex-based differences are consistent with previous studies [41], which demonstrated that female mice possessed more resident immunomodulatory CD4^+^ T cells and mounted a more efficient immune response in acute inflammation. Notably, it has also been shown that biological sex, among other variables, significantly influenced mouse mortality outcomes across 14 days [42]. The combination of this data leads to the conclusion that sex-based divergence in sepsis outcomes may begin at the transcriptional level well before it becomes clinically or physiologically evident.

Although our study used mice, we can tentatively extrapolate these findings to the clinical context specifically as it relates to sex-dependent differences, which have been documented in the clinical literature. For instance, there is evidence that men may have a higher sepsis incidence [10,43], but that women have an 8% higher rate of in-hospital mortality due to sepsis [44]. It has also been reported that women exhibit lower long-term mortality and fewer ICU admissions compared to men [9,10,12,45]. Previous clinical studies have also suggested potential sex-based differences in septic renal and respiratory injuries, although the findings thus far are inconclusive [46,47].

These sex-based differences may reflect the complex interplay between sex hormones, immune cell composition, and inflammatory regulation. For example, estrogen has been shown to enhance innate immune responses while promoting resolution of inflammation, which may contribute to both early elevated risk and long-term resilience [10,48]. Additionally, sex-based differences in adaptive immunity – including higher proportions of regulatory T cells and immunomodulatory CD4^+^ lymphocytes in females – may provide enhanced control of systemic inflammation and better outcomes following sepsis [41,47]. Consistent with these observations, our transcriptomic findings revealed stronger early activation of innate immune pathways in females.

Taken together, it is plausible that females may have an advantage in mounting a more robust and efficient immune response to infection compared to males, which, in the initial stages of sepsis, is characterized by stronger inflammatory activation. While this heightened and dysregulated immunomodulatory response may initially lead to more tissue and organ damage and even elevated mortality, in the long term, this response may favour female survival over that of males [9,49]. This speculation emphasizes the importance of future studies to elucidate the impact of early transcriptomic and other pathophysiological differences between males and females, as it relates to long-term clinical outcomes, such as hospitalization, disease progression, and ultimately, mortality.

While our study provides insights into organ and sex-specific transcriptional responses during sepsis, several limitations must be acknowledged. Although liver tissue was included in the combined population analysis, limited sample size precluded sex-stratified analysis of hepatic transcriptional responses. Furthermore, this study focused exclusively on transcriptional outcomes at a single early time point, and protein-level validations or longitudinal assessments were not performed. Nevertheless, our study demonstrates that biological sex differences in the septic response can be detected in early acute sepsis without apparent physiological differences. Furthermore, our study provides a critical, initial framework that can be expanded to include additional biological variables, such as age, diet, and prior exercise training, among others [20,47,50–53]. Whereas the current study emphasized organ and sex specific differences, studies including these would further advance the development of preclinical models that more accurately reflect septic pathophysiology in human patients. Such studies are needed to provide a stronger foundation supporting future personalized medicine approaches to reduce the burden of sepsis in patients.

## Acknowledgements

The authors thank David Carter at London Regional Genomics Centre for help with the alignment and initial RNAseq analysis. Thanks to the numerous undergraduate students that assisted with experimental procedures and data entry.

## Supplementary Materials

**Sup Fig S1. GSEA in the Kidney during Sepsis in Male vs. Female Mice.**

Within the kidney, males exhibited enrichment of most **(A)** Proliferation & Cell Cycle and **(B)** Metabolism pathways under dextrose (control) conditions, with further enhancement of these pathways under slurry (septic) conditions. Females and males showed minor enrichment of **(C)** Canonical Signalling pathways under dextrose (control) conditions, with further enhancement under slurry (septic) conditions. Females exhibited stronger enrichment than males in **(D)** Development & Differentiation pathways under both dextrose (control) and slurry (septic) conditions. The cutoff for significance is FDR ≤ 0.1. The normalized enrichment score (NES) reflects the direction and magnitude of pathway enrichment in females relative to males. The gene count per pathway is identified by the size of the circle.

**Sup Fig S2. GSEA in the Lung during Sepsis in Male vs. Female Mice.**

Within the lung, males exhibited enrichment in most of the pathways – **(A)** Proliferation & Cell Cycle, **(B)** Metabolism, **(C)** Canonical Signalling, and **(D)** Development & Differentiation – under dextrose (control) conditions, while maintaining enrichment in **(A)** Proliferation & Cell Cycle and **(B)** Metabolism under slurry (septic) conditions. Females showed no significant enrichment in any pathways under dextrose (control) conditions, while exhibiting significant enrichment in subsets of the **(C)** Canonical Signalling and **(D)** Development & Differentiation pathways under slurry (septic) conditions. The cutoff for significance is FDR ≤ 0.1. The normalized enrichment score (NES) reflects the direction and magnitude of pathway enrichment in females relative to males. The gene count per pathway is identified by the size of the circle.

## Notes

### Competing Interest Statement

The authors have declared no competing interest.

